# Sex differences in durability following heavy intensity cycling exercise in trained athletes

**DOI:** 10.1101/2025.08.26.672326

**Authors:** Elisa Pastorio, Padraig Spillane, Emma Squires, Leila Benyahia, Hannah K Wilson, Patrick Swain, Marta Colosio, Chiara Felles, Andrea Menditto, Samuel Clarke, Beth Minion, Will Pearmain, Callum G Brownstein, Simone Porcelli, Paul Ansdell

## Abstract

The ability to withstand impairments in key physiological variables during prolonged exercise, known as ‘durability’, is emerging as an important factor in cycling performance. While females possess physiological characteristics that could confer enhanced durability relative to males, little is known about potential sex differences.

32 trained cyclists (16 males and 16 females) performed an incremental exercise test to exhaustion in visit 1. In visit 2 they performed 90 minutes of heavy intensity cycling (HVY) at 110% of gas exchange threshold (GET), followed by another incremental test. During HVY, pulmonary gas exchange (V̇O_2_ and V̇CO_2_) ventilation (V̇E), heart rate (HR), rating of perceived exertion (RPE), near-infrared spectroscopy and electromyography were recorded, and blood lactate (BLa) was collected. Before and after HVY, maximal voluntary contraction (MVIC), voluntary activation (VA) and potentiated twitches (100Hz, 10Hz, Q_tw·pot_) of the knee extensors were assessed.

Power at GET (-16±15% vs -2±13%) and respiratory compensation point (-13±10% vs -6±9%) decreased more in males than females (P≤0.049). All aspects of neuromuscular function decreased from pre to post (all P<0.001), without sex differences (P≥0.096). During HVY, HR, V̇O_2_ (%peak), relative energy expenditure increased more in males (P≤0.008), whereas respiratory exchange ratio decreased more in females (P=0.001). BLa was higher in males than females (P=0.030). Muscle oxygen extraction was lower (P=0.004) and tissue saturation index higher for females (P<0.001).

The smaller reductions exhibited by females in submaximal thresholds, associated with lesser derangements to oxidative efficiency, highlight the need to consider sex-specific training prescription and pacing strategies for long duration events.

**Key Points:**

- Durability, as measured by the reduction in incremental exercise test outcomes, is relatively unexplored in females compared to males, despite physiological sex differences that might confer a female advantage.
- After 90 minutes of heavy intensity cycling, males demonstrated greater reductions in the power outputs associated with gas exchange threshold and respiratory compensation point.
- The maximal rate of oxygen consumption and incremental test peak power output decreased similarly in both sexes.
- These changes are associated with greater carbohydrate metabolism and losses of efficiency in males, whereas no sex differences were observed in neuromuscular fatigue.

## INTRODUCTION

The power that athletes can sustain during endurance events is closely associated with the maximal O_2_ uptake (V̇O_2max_), the fractional utilisation of V̇O_2max_ at lactate threshold, and the economy during sub-maximal exercise (Joyner & Coyle, 2008). These variables are typically assessed in fresh conditions, however, recent studies (Clark et al. 2018, 2019a, 2019b; Stevenson et al. 2022) have showed that these physiological pillars of endurance performance are not fixed but change over time due to their progressive deterioration as exercise proceeds. For instance, Clark et al. (2018, 2019a, 2019b) showed that after 2 hours of cycling in the heavy intensity domain, both the critical power (CP) and the fixed amount of available energy above CP (W’) were reduced by 8-11% and 17-22%, respectively. Moreover, Stevenson et al. (2022) demonstrated that 2 hours of moderate intensity cycle exercise reduces lactate threshold and gas exchange threshold (GET) by 10%. The ability of an athlete to mitigate this decrease in metabolic threshold(s) has recently been termed “resilience” or “durability” (Maunder et al. 2021; Jones, 2024) and it has been proposed as a new marker of elite performance, given its important performance implications (Van Erp et al. 2021). In this regard, a recent study by Hamilton et al. (2024) showed that durability of the moderate-to-heavy-intensity transition (VT_1_) is an important performance parameter, as more durable athletes exhibited smaller reductions in 5-min time trial performance following 150 minutes of moderate intensity cycling.

In addition to lab-based studies, recent field-based evidence (Van Erp et al. 2021; Gallo et al. 2022; Mateo-March et al. 2022) suggests that the capability of a rider to maintain power following previous accumulated work might be a characteristic of successful competitive male road cyclists. Indeed, Gallo et al. (2022) found that the superior durability, measured as decrement in the mean power produced during maximal efforts of 1,5 and 20 min after the accumulation of up to 50 kJ/kg of work, was the only parameter able to differentiate professionals from under-23 riders. Similarly, Van Erp et al. (2021) reported that higher standard cyclists demonstrated greater ability to maintain high mean power outputs over fixed durations following the accumulation of up to 50 kJ/kg work done compared to category 2 cyclists, despite similar record power profiles in fresh condition. Overall, this recent evidence suggests that, during cycling competitions, a rider’s capacity to express high power outputs following a substantial amount of work done (such as after hours of racing on road, or at the end of exhausting high-intensity competitions on track) represents a key determinant of success in professional cycling (Jones & Vanhatalo, 2017; Van Erp et al. 2021; Maunder et al. 2022). Therefore, it is important for competitive cyclists to assess their performance both in fresh state and after fatiguing cycling tasks, in order to quantify their “durability” and ultimately their performance.

Currently, the physiological determinants of durability are not completely understood. However, the ability to delay the onset and reduce the magnitude of performance fatigability (i.e., decrements in neuromuscular function) during prolonged exercise is essential to prevent the loss of efficiency (i.e., the rise in energy cost for a given work rate, also defined “slow component” of V̇O_2_; Grassi et al. 2015) that occurs over time (Jones et al. 2011). This loss of efficiency is typically attributed to changes in motor unit recruitment profiles (i.e. greater recruitment of type II fibres owing to impaired contractile function in already recruited fibres) and substrate utilisation (i.e. muscle glycogen depletion, greater fat oxidation; Gollnick et al. 1973; Vøllestad & Blom, 1985). In this regard, it is well-known that females are more fatigue-resistant than males during various tasks when assessed with measures of neuromuscular function (Hunter, 2016). Indeed, both after two hours of constant-load cycling at the first ventilatory threshold (Glace et al. 2013) and following constant-load cycling in the heavy and severe intensity domains (Ansdell et al. 2020; Azevedo et al. 2021) females experienced a greater preservation of contractile function of the knee extensors than males. These findings suggest that some sex-specific anatomical and physiological features of females, such as the greater fat oxidation and better preservation of glycogen stores (Tarnopolsky, 2008), higher proportion of type I fibres (Roepstorff et al. 2006), and greater muscle perfusion (Parker et al. 2007; Hammer et al. 2023), as well as the superior muscle oxidative capacity (Cardinale et al. 2018), might contribute to delay the onset of fatigue in females during prolonged endurance events (Temesi et al. 2015; Besson et al. 2022). What remains unexplored is whether these physiological sex differences promote superior durability in females, allowing them to better maintain performance compared to males during prolonged events. The importance of this is underscored by the rapid increase in the participation of females in cycling (ProCyclingStats, 2025), as well as in the recognition of female professional cyclists with the introduction of Women’s World Tour Series in 2016 (Van Erp, 2019). Accordingly, there is an urgent need to increase knowledge through performance-based research in females and consequently develop more sex-specific training guidelines (Costello et al. 2014; Emmonds et al. 2019; McNulty et al. 2020; Cowley et al. 2021). To date, most studies have investigated durability in male athletes and only one recent field based-study included also female cyclists (Mateo March et al. 2025), reporting similar relative record power profile reductions after >10 kJ/kg, but higher relative decay in females after 20 kJ/kg for 1-min, 5-min, and 20-min efforts. However, the authors acknowledged differences in training history, as well as in the physiological demands of training and competition between their cohorts of males and females, which could confound their findings.

This study aimed to investigate durability, and its physiological underpinnings, in well-trained male and female cyclists following 90 minutes of constant-load cycling in the heavy intensity domain. It was hypothesised that (i) females would have a better durability than males, as measured by the decrease in maximal and submaximal markers of performance; (ii) females would experience a slower derangement in cardiopulmonary, metabolic, and neuromuscular parameters during the 90 minutes of heavy intensity exercise.

## METHODS

### Ethical approval

Participants were fully informed of the aim and procedures of the study, including its risks and benefits, before signing the written informed consent form. All procedures conformed to the Declaration of Helsinki and were approved by the institutional ethical approval from the Northumbria University Health and Life Sciences Research Ethics Committee (submission reference: 2024-6525-6208)

### Participants

Using a small effect size (ηp^2^ = 0.022) for a potential sex difference in the change in the power at ventilatory threshold (reliability statistic r = 0.90; Pallerés et al. 2016), with the parameters: α = 0.05, 1-β = 0.8, the *a priori* sample size calculation revealed that a minimum of 32 participants were required. Therefore, 32 healthy competitive cyclists and triathletes, including 16 females (mean ± SD age: 27 ± 4 yr; stature: 164 ± 6 cm; mass: 56.8 ± 4.4 kg; V̇O_2peak_: 51 ± 3 ml · kg^-1^ · min^-1^) and 16 males (age: 29 ± 6 yr; stature: 181 ± 7 cm; mass: 75.9 ± 10.0 kg; V̇O_2peak_: 58 ± 6 ml · kg^-1^ · min^-1^) were enrolled in the study after providing written informed consent. All participants were well-trained (Performance Level 3 or above, De Pauw et al. 2013; Decroix et al. 2016), competed in cycling or triathlon races and were familiar with ≥ 2 hours exercise bouts. To ensure that the cardiovascular fitness of the two groups was similar, participants had to have a minimum V̇O_2peak_ (determined in the initial visit, see below) of 48 ml · kg^-1^· min^-1^ for females and 55 ml · kg-^1^ · min^-1^ for males. These values correspond to equitable performance levels in sports science research (De Pauw et al. 2013; Decroix et al. 2016). Two additional participants were recruited, but they did not complete the experimental visit due to not meeting the inclusion criteria. Moreover, in order to be included in the study, all participants had to be aged between 18 and 40 years old, had to be free from cardiovascular, respiratory, or neurological diseases or illness as well as musculoskeletal injuries that could affect cycling performance in the last 6 months. Regarding their training characteristics, participants reported training 5±2 days/week and accumulating 10±4 hours of training/week, without sex differences (*P ≥* 0.182).

This was a multi-site study, where participants were recruited and tested at Northumbria University (n=11), Newcastle University (n=12) and at the University of Pavia (n=9) using the same methods and protocols.

#### Female participants

For female participants, both naturally cycling females and hormonal contraceptive users were included. More specifically, naturally cycling participants reporting a regular menstrual cycle duration (>21 and <35 days), and not been using hormonal contraceptives for at least 6 months, had no MC-related irregularities (e.g., amenorrhea), or conditions (e.g., polycystic ovarian syndrome, endometriosis, and pregnancy) known to affect the hypothalamic–pituitary–ovarian axis (McNulty et al. 2023) were included. Moreover, females who were taking a combined monophasic oral contraceptive pill (OCP) (containing a dose of 30 *µ*g of EE) for at least 6 months were included, as well as females taking any other kind of continuous hormonal contraceptives (implants, intrauterine devices/coil, etc). In our female cohort, n=11 were naturally cycling females, n=1 was taking a combined monophasic pill, n=2 were using a Mirena coil, n=1 reported to have a Nexplanon contraceptive implant (progesterone only) and n=1 had a vaginal ring.

Naturally cycling females were tested during the early follicular phase of the menstrual cycle (both low oestrogen and low progesterone levels) whereas the oral contraceptive and the vaginal ring users were tested during the pill withdrawal phase and the removal period, respectively. During these time points, the hormonal environments are expected to be similar between these groups (Elliott-Sale et al. 2021). The other females reporting to have the Mirena coil or the Nexplanon contraceptive implant, which are progesterone only, were not tested during a specific phase as their hormonal environment remained constant during usage.

### Experimental design

Participants visited the laboratory on two occasions, separated by at least 72 hours of rest, and at the same time of day. The first visit involved a familiarisation with neuromuscular measures, followed by a cycling incremental exercise test to the limit of tolerance. In the second visit participants performed fatiguing cycling exercise, involving 90 minutes of heavy intensity cycling at 110% of GET (Brownstein et al. 2022) followed by the same incremental test performed in visit 1. Before and immediately (commencing within 60 seconds) after the fatiguing exercise, maximal voluntary isometric contraction force (MVIC), voluntary activation (VA) and potentiated twitches (high-frequency 100 Hz doublet, Db_100_; low-frequency 10 Hz doublet, Db_10_; single stimulation, Q_tw·pot_) of the quadriceps were assessed. During both incremental exercise tests, pulmonary gas exchange, heart rate (HR), and rating of perceived exertion (RPE) were recorded, and blood lactate (BLa) sampling were collected. In addition to these measurements, during the 90 minutes of heavy intensity cycling, near-infrared spectroscopy (NIRS), and electromyography were also recorded.

All participants were asked to maintain their habitual diet throughout the study, recorded their diet 24 h prior to the first lab visit and replicated the same diet prior to the second visit, and were instructed to eat a high carbohydrate meal before attending each visit. Participants arrived at the lab ≥ 2h post prandial and in a hydrated state. They refrained from alcohol and vigorous exercise < 48h prior to visits, consumption of nutritional supplements/ergogenic aids (e.g. sodium bicarbonate, nitrates) < 24 h prior to visits and consumption of caffeine < 6h prior to each visit.

#### Visit 1: Familiarisation and Incremental Test

This visit began with participants being familiarised with the neuromuscular stimulation techniques. After the determination of the femoral nerve stimulation intensity, then the participants performed a series of warm up contractions increasing from 50% perceived effort to 90%. Hereafter, a neuromuscular assessment was performed (see full details below), before participants moved to the cycle ergometer (Lode Excalibur, Lode B.V., Groningen Country, NL) to perform a modified step-ramp protocol (Brownstein et al. 2022), from that developed by Iannetta et al. (2019).

Resting measurements of pulmonary gas exchange and heart rate were recorded for three minutes, and a blood lactate sample was taken. Then, the step-ramp protocol began with two minutes of cycling at 20 W, followed by six minutes of cycling at 1.5x body mass (males) or 1.3x body mass (females). Thereafter, participants completed four minutes of cycling at 20 W before commencing the ramp test, which increased the work rate by 20 W· min^-1^ for females, or 25 W· min^-1^ for males. The ramp test continued until task failure, defined as the point at which cadence decreased by 10 rpm from the participant’s self-selected cadence for more than 5-s despite strong verbal encouragement.

#### Visit 2: Fatiguing Task and Durability assessment

This visit began with a neuromuscular assessment, before participants moved to the cycle ergometer and provided a resting blood lactate sample, and recordings of pulmonary gas exchange, near-infrared spectroscopy, and electromyography were taken.

Participants began with a warm up consisting of a 5-minute stage at 75% of the power output associated with GET, after which the power output instantly increased to the target power output for the 90 minutes of heavy-intensity exercise at 110% GET. Participants were instructed to maintain their preferred cadence (between a range of 80 and 90 rpm) and they were given the choice of listening to music during the 90 minutes trial and to have a visible clock showing the time remaining. Participants were also allowed to drink water ad libitum but not consume any food or calorie-containing drinks during the test. Immediately (commencing within 60 seconds) after the completion of the fatiguing task, participants repeated the neuromuscular assessments and then performed the same step-ramp protocol as the first visit. Following the step-ramp protocol, participants laid supine, whilst blood flow was occluded for 5 minutes to allow a physiological calibration of NIRS signals (see below).

During the fatiguing task, heart rate, near-infrared spectroscopy and electromyography were recorded continuously, whereas breath-by-breath gas exchange measures were taken for the first 15 minutes, and then for 5 minutes, every 15 minutes thereafter. Ratings of perceived exertion (6-20 scale, Borg, 1982) and blood lactate samples were collected every 15 minutes. The full protocol is displayed in Figure 1.

**Figure 1.**
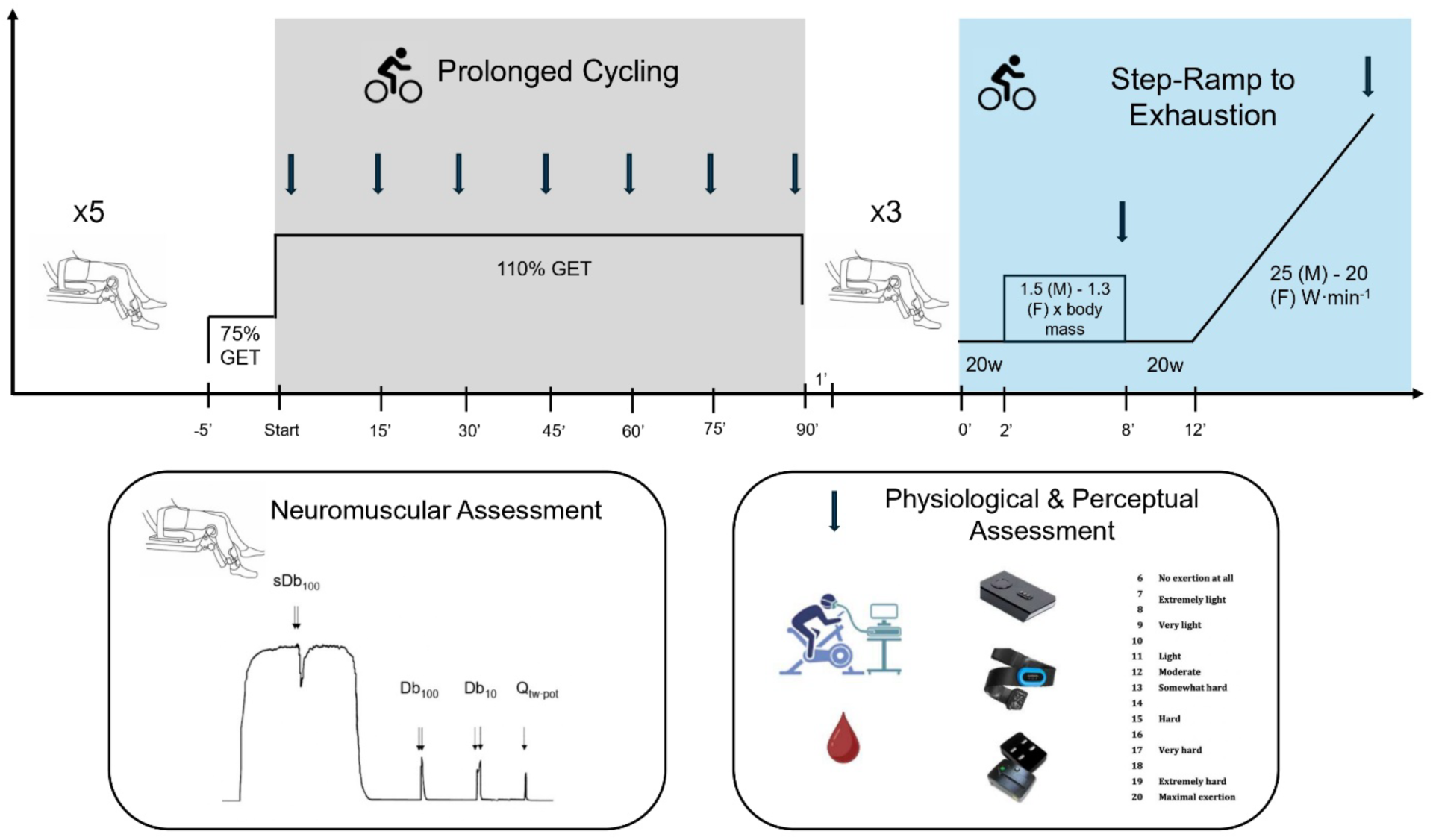
Graphical representation of the experimental procedure performed in visit 2. The y axis represents the exercise intensity during each stage of the protocol whereas the x axis shows the timeline of the protocol. The chair figure represents a neuromuscular assessment (consisting of MVICs and potentiated twitches at rest). The colored background highlights the cycling stages, whereas the arrows indicate time points for physiological and perceptual assessment (breath by breath gas exchanges, blood lactate, heart rate, electromyography and rate of perceived exertion).

### Experimental techniques

#### Pulmonary Gas Exchange

During all visits, pulmonary gas exchange and ventilation were measured breath-by-breath using an online system (Vyntus CPX, Jaeger, CareFusion, Germany). With minute ventilation (V̇E), oxygen consumption (V̇O_2_), carbon dioxide production (V̇CO_2_) and respiratory exchange ratio (RER) were quantified. Before each test, the analysers for oxygen and carbon dioxide were calibrated using ambient air and a gas of known concentration (15% O_2_ and 4.97% CO_2_). Ventilatory volumes were calibrated using a digital turbine transducer at high (2 L s^-1^) and low (0.2 L s^-1^) flow rates.

#### Neuromuscular assessments

Following warm up contractions, baseline neuromuscular assessments consisted of five MVICs separated by 60s. During the final three MVICs, a high-frequency doublet electrical stimulation [100 Hz, 10-ms interstimulus interval (sDb_100_)] was superimposed to the contraction at peak force. Subsequently, once the muscle was relaxed, a series of potentiated high (100Hz) and low-frequency (10 Hz, 100-ms interstimulus interval, Db_10_) doublet stimulations, as well as a single twitch (Q_tw·pot_), were delivered. This assessment protocol was used to quantify MVIC force, voluntary activation (VA), and contractile function through Db_100,_ Db_10:100_ and Q_tw.pot_. At the end of the fatiguing task, in the interest of time the neuromuscular assessment was repeated but with three MVICs instead of five, as re-potentiation of twitches was not required (Kufel et al. 2002).

#### Femoral Nerve Stimulation

Electrical stimuli (200 µs duration) were delivered to the femoral nerve via 32 mm-diameter surface electrodes (CF3200; Nidd Valley Medical, North Yorkshire, UK) using a constant-current stimulator (DS7AH, Digitimer, Welwyn Garden City, Hertfordshire, UK). The cathode was placed high in the femoral triangle over the nerve, and the anode was positioned midway between the greater trochanter and the iliac crest. The cathode was repositioned, if necessary, to the point that elicits the largest quadriceps twitch amplitude (Q_tw_). Stimulations began at 20 mA and increased by 20 mA until a plateau in Q_tw_ occurred. This value was then increased by 30% to ensure supramaximal stimulation was delivered during the protocol. Mean stimulus intensity was not different between males (309 ±108) and females (260 ±152 mA, *P =* 0.360).

#### Force and Electromyography

For the neuromuscular assessments, participants were seated on a custom-built chair, with force (N) measured using a calibrated load cell (MuscleLab force sensor 300, Ergotest technology, Norway or SML load cell, Interface, Scottsdale, AZ). The load cell was attached to the participant’s dominant leg, 2 cm superior to the ankle malleoli, using a non-compliant cuff. The load cell height was adjusted to ensure a direct line with the applied force for each participant. Participants sat upright with knee and hip angles kept at 90° flexion. The chair was placed ∼1 m adjacent to the cycle ergometer to facilitate movement and save time for the post assessment. Force was sampled continuously (1000 Hz), and acquired for offline analysis (Spike 2, Cambridge Electronic Design, Cambridge, UK or LabChart 8, ADInstruments, Bella Vista, Australia).

EMG signals were recorded continuously at 2000Hz throughout both visits using wireless sensors (10 mm inter-electrode distance; Trigno Avanti, Delsys, MA, USA or Pico EMG, Cometa, Bareggio, Italy). Sensors were placed over the participant’s dominant leg on Rectus Femoris (RF), Vastus Lateralis (VL) and Vastus Medialis (VM) consistent with SENIAM guidelines (Hermens et al. 2000). Prior to placement, the skin-electrode contact area was shaved, abraded and cleaned using a 70% IPA alcohol wipe (FastAid, Robinson Healthcare, Workshop, UK). The raw EMG signals were amplified (gain x 150) and, using a custom script (MATLAB 2024b, MathsWorks Inc., Natick, MA, US), filtered using a 20-450 Hz 4^th^ order Butterworth bandpass filter.

Signals were amplified: gain × 150 for EMG (Delsys Trigno EMG systems, Boston, MA, USA or Pico EMG, Cometa, Bareggio, Italy) and × 300 for force (CED 1902; Cambridge Electronic Design, Cambridge, UK or PowerLab 8/35, ADIntruments), bandpass filtered (EMG only: 20-450 Hz), digitized (EMG: 2 kHz; force: 5 kHz; CED 1401, Cambridge Electronic Design or PowerLab 8/35, ADInstruments or Load Cell Adapter, Delsys, Natick, MA), and analysed offline (force: Spike2 v8, Cambridge Electronic Design, EMG: R2024; Mathworks Inc., Natick, MA)

#### Near Infrared Spectroscopy

A wireless, portable, and continuous-wave spatially resolved NIRS light photometer (Portamon, Artinis Medical Systems) was used to evaluate relative changes in oxy – (O_2_HbMb) and deoxy-(HHbMb) heamoglobin and myoglobin, as well as tissue saturation index (TSI: [O_2_Hb Mb]/total [Hb+Mb] × 100), sampled at 10 Hz. The probe was placed on the lower third of vastus lateralis muscle (∼ 10 cm above the knee joint). The probe was held in place by an elasticised, tensor bandage and covered by an opaque, dark material to avoid motion and ambient light influences (Pilotto et al. 2022). Before the starting of both the incremental test and the fatiguing task, baseline TSI, O_2_HbMb and HHbMb were recorded over 3 minutes of rest. At the end of each visit, a prolonged ischemia (5 min) from femoral artery occlusion by cuff inflation with pressure above 120% of limb occlusion pressure, which was not different between sexes (*P=* 0.271), was performed by using a 13×85 cm rapid-inflation pressure cuff (Delfi Medical Innovations Inc., Vancouver BC, Canada or E20, or Hokanson, Place Bellevue, WA USA) placed proximally on the same thigh and attached to a cuff-inflator. This procedure was used to perform the physiological calibration of the NIRS signal and the relative changes in VL O_2_HbMb and HHbMb during the fatiguing task were expressed as percentage change from the minimum (0%) and the maximum (100%) value recorded during the physiological calibration.

#### Blood Lactate Sampling

Blood lactate was sampled via capillary puncture technique with a 10 μl sample taken from the earlobe of each participant. Samples were immediately analysed for the concentration of lactate (mmol·L^-1^, Biosen, EKF Diagnostics) and used to assess changes during the fatiguing task.

### Data Analysis

All MVICs were recorded, and the single contraction with the greatest peak force (250-ms window before the superimposed electrical stimulation) (Merletti & Farina, 2016) used for further analyses. To quantify impairments to the central nervous system drive, knee extensors voluntary activation (VA) was calculated using the interpolated twitch technique and calculated from the equation (Allen et al. 1995): VA (%) = (1 – [sDb_100_/ Db_100_] × 100), where sDb_100_ is the amplitude of the high-frequency doublet superimposed twitch force measured during MVIC, and Db_100_ is the amplitude of the high frequency doublet twitch force assessed 2 s post-MVC on relaxed muscle. Changes in skeletal muscle contractile properties were assessed by variations in the amplitudes of Q_tw·pot_, Db_100_, and Db_10:100_ (Jones, 1996). Peak-to-peak EMG amplitude elicited by single femoral nerve electrical stimulation was measured to assess sarcolemma excitability (M_max_). The filtered EMG data were then root-mean-squared across a 200 ms moving window (EMG_RMS_). The maximum EMG_RMS_ elicited during the MVIC was expressed as a percentage of M_max_. For measurements of EMG during the incremental and fatiguing cycling tasks, the peak EMG_RMS_ during each EMG burst (i.e., each VL, VM, and RF contraction per pedal rotation) was automatically identified and expressed as a percentage of both the pre-exercise maximum EMG_RMS_ as well as the pre-exercise M_max_ using a custom-built script (R2024; Mathworks Inc., Natick, MA). Cycling EMG_RMS_ was averaged at the same time points as gas exchange measurements and NIRS data.

The rate of V̇O_2_, V̇CO_2_, and V̇E were recorded during both visits and were exported in 5 seconds intervals. Gas exchange threshold (GET) and respiratory compensation point (RCP) were visually, individually, and independently determined by two expert investigators after they agreed on the value set, using multiple gas exchange and ventilatory equivalent criteria through consideration of V̇O_2_, V̇CO_2_, end-tidal partial pressures for CO_2_ and O_2_, and V̇E/V̇CO_2_ and V̇E/O_2_ (Wasserman et al. 1973; Beaver et al., 1986). The power output associated with the GET was determined according to the methods outlined by Brownstein et al. (2022), by linearly interpolating the V̇O_2_ versus power output relationship after the V̇O_2_ was left-shifted by a time interval corresponding to the MRT. Using the mean response time (MRT) from the moderate intensity step, the power output associated with GET was left-shifted to account for the time delay between changes in muscle metabolism being reflected in pulmonary gas exchange (Iannetta et al. 2019). V̇O_2peak_ was defined as the highest 30 s rolling average of V̇O_2_, and peak power output (P_max_) was considered as the last power output that each participant was able to complete during the ramp test before task failure.

During the 90 minutes of fatiguing cycling, the variables as V̇O_2_, V̇CO_2_, V̇E, RER as well as O_2_HbMb and HHbMb, were averaged from 4 to 6 minutes after exercise commencement (in order to avoid phase I and II onset kinetics) and during the last 2 minutes of each 5 minutes time intervals (13 to 15 min, 28 to 30 min, 43 to 45 min, 58 to 60 min, 73 to 75 min, 88 to 90 min). Relative whole-body energy expenditure (EE/kg) expressed in J·s·kg^-1^ was calculated according to the equation (x) = [(281.67 × V̇̇O_2_) (L·min^-1^) + (80.65 × ̇ V̇CO_2_) (L·min^-1^) /kg (Péronnet & Massicotte, 1991). Gross efficiency (GE) (%) was calculated as Work rate (W)/ Energy Expenditure (J·s) × 100.

### Statistical Analysis

Results are presented as means ± SD within the text and figures. Statistical significance was set at an α level of 0.05. Normality of the data was assessed by the Shapiro Wilk test, with no data requiring transformation. Assumptions of sphericity were explored and controlled for all variables with the Greenhouse–Geisser adjustment, where necessary. Sex comparisons for P_max_, V̇O_2peak_, V̇CO_2peak_, V̇E_peak_, HR_peak ,_ Bla_peak ,_ PO at GET and at RCP, GET and RCP (%V̇O_2peak_) and GET and RCP (%HR_max_) were assessed with independent samples t tests. Exercise-induced changes in MVIC, VA, Db_100_, Db_10:100_, Q_tw·pot_, PO at GET, PO at RCP, P_max_, V̇O_2peak,_ BLa, HR, V̇O_2_, V̇CO_2_, V̇E, RER, EE, GE, EMG_RMS_, TSI%, O_2_HbMb, HHbMb and RPE were assessed with a two-way repeated measures ANOVA. Each ANOVAs involved sex and time as the independent variables. Significant sex and time main effects or sex × time interaction effects were further explored using Bonferroni-corrected pairwise comparisons. Partial eta squared (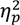) was calculated to estimate effect sizes, with values representing small (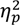 < 0.13), medium (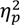≥0.13, <0.26) and large (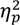≥0.26) effects. The statistical software package Prism 10 (GraphPad, Software, Inc., San Diego, CA) was used to analyze data.

## RESULTS

### Baseline incremental test results

The variables recorded during the incremental ramp test performed in visit 1 are displayed in Table 1. As expected, males showed greater values both for absolute and relative V̇O_2peak_, P_max_, V̇CO_2peak_, V̇E_peak_, power outputs associated at GET and RCP (all *P* < 0.001). HR_peak_, Bla_peak_, V̇O_2_ (%peak) and HR (%peak) associated at GET and RCP were not different between males and females (*P* ≥ 0.053). The average performance level (De Pauw et al. 2013; Decroix et al. 2016) was similar between males and females (3 ± 1 vs 3 ± 1, *P* = 0.265).

As expected, before HYV, males showed greater MVIC (630 ± 148 vs 378 ± 75 N, *P <* 0.001), resting twitch responses for Db_100_ (248 ± 81 vs 170 ± 51 N, *P =* 0.005), Db_10_ (247 ± 76 vs 164 ± 44 N, *P =* 0.001), Q_tw·pot_ (168 ± 58 vs 108 ± 29 N, *P =* 0.001) and M_max_ (11.15 ± 6.05 vs 6.94 ± 4.54 mV, *P =* 0.042) than females. VA was not different between sexes (94 ± 4 vs 91 ± 5 %, *P =* 0.136).

**Table 1.** Participant demographics and baseline incremental test results in males and females. Values are presented as mean ± SD.

|  | Males | Females | P value |
| --- | --- | --- | --- |
| N | 16 | 16 | n/a |
| Age (years) | $29 \pm 6$ | $27 \pm 4$ | 0.431 |
| Stature (cm) | $181 \pm 7$ | $164 \pm 6$ | **< 0.001** |
| Mass (Kg) | $75.9 \pm 10$ | $56.8 \pm 4.4$ | **< 0.001** |
| Training (h·week<sup>-1</sup>) | $9 \pm 4$ | $11 \pm 3$ | 0.335 |
| **Incremental test** |  |  |  |
| $\dot{V}O_{2peak}$ (L·min<sup>-1</sup>) | $4.40 \pm 0.60$ | $2.91 \pm 0.35$ | **< 0.001** |
| $\dot{V}O_{2peak}$ (ml·kg<sup>-1</sup>·min<sup>-1</sup>) | $58.3 \pm 6.8$ | $51.2 \pm 3.1$ | **0.001** |
| $P_{max}$ (W) | $410 \pm 56$ | $270 \pm 35$ | **< 0.001** |
| $P_{max}$ (W·kg<sup>-1</sup>) | $5.4 \pm 0.7$ | $4.7 \pm 0.4$ | **0.001** |
| $\dot{V}CO_{2peak}$ (L·min<sup>-1</sup>) | $5.32 \pm 0.51$ | $3.47 \pm 0.46$ | **< 0.001** |
| $\dot{V}E_{peak}$ (L·min<sup>-1</sup>) | $183.95 \pm 18.45$ | $118.19 \pm 9.25$ | **< 0.001** |
| $HR_{peak}$ (bpm) | $185 \pm 17$ | $184 \pm 10$ | 0.142 |
| $Bla_{peak}$ (mmol<sup>-1</sup>) | $10.2 \pm 3.1$ | $9.9 \pm 1.7$ | 0.681 |
| GET (W) | $226 \pm 42$ | $135 \pm 21$ | **< 0.001** |
| GET (%$\dot{V}O_{2peak}$) | $66 \pm 4\%$ | $63 \pm 5\%$ | 0.053 |
| GET (%$HR_{max}$) | $78 \pm 4\%$ | $78 \pm 5\%$ | 0.749 |
| RCP (W) | $331 \pm 25$ | $222 \pm 35$ | **< 0.001** |
| RCP (%$\dot{V}O_{2peak}$) | $89 \pm 3\%$ | $89 \pm 5\%$ | 0.607 |
| RCP (%$HR_{max}$) | $92 \pm 3\%$ | $93 \pm 4\%$ | 0.561 |

Abbreviation: $\dot{V}O_{2peak}$ , peak oxygen uptake; $P_{max}$ , maximal power; $\dot{V}CO_{2peak}$ , peak carbo dioxide production; $\dot{V}E_{peak}$ , peak ventilation; $HR_{peak}$ , peak heart rate; $Bla_{peak}$ , peak blood lactate; GET, gas exchange threshold; RCP, respiratory compensation point.

### Change in incremental test results after heavy intensity cycling

#### Constant heavy intensity cycling

Males performed HVY at a greater absolute intensity than females (247 ± 48 vs 149 ± 23 W, *P* < 0.001) but at the same relative intensity, which corresponded to 110% of power output associated at GET. Indeed, at the beginning of the fatiguing cycling trial (4-6 min) the V̇O_2_ (%V̇O_2GET_) at the target intensity was not different between males (112.7 ± 7.4%V̇O_2GET_) and females (112.7 ± 10.3%V̇O_2GET_, *P* = 0.982). Of the whole sample of 32 participants, three male participants reached task failure at 60, 78 and 86 minutes of exercise and one female participant terminated the trial at 60 minutes of cycling. However, despite achieving task failure, they all completed the ramp test POST.

#### Incremental ramp test POST

Changes in incremental test results after HVY are displayed in Figure 2. A time effect was found for P_max_ (-10 ± 8%; F _1,30_ = 51.83, *P* < 0.001, 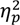 = 0.63; Figure 2E), for V̇O_2peak_ (-7 ± 7%; F _1,59_ = 27.63, *P* < 0.001, 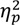 = 0.32; Figure 2F), V̇CO_2peak_ (-15 ± 12%; F _1,59_ = 50.84, *P* < 0.001, 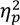 = 0.46), BLa_peak_ (-45 ± 20%; F _1,59_ = 161.0, *P* < 0.001, 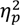 = 0.73), and RER_peak_ (-9 ± 8%; F _1,59_ = 39.8, *P* < 0.001, 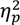 = 0.40), with no effects of sex, or time x sex interactions (*P* ≥ 0.072). A time effect (F _1,59_ = 28.63, *P* <0.001, 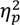 = 0.33), sex effect (F _1,59_ = 4.04, *P* = 0.049, 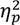 = 0.06) and a time x sex interaction (F _1,59_ = 4.04, *P* = 0.049, 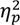 = 0.06) was found for V̇E_peak_, which decreased more in males compared to females (-18 ± 16% vs -9 ± 10%, *P* = 0.049).

A time effect was found for PO at GET (F_1,30_ = 13.6, *P* ≤ 0.001, 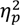 0.31; Figure 2A) and at RCP (F_1,30_ = 29.89, *P* ≤ 0.001, 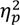 = 0.50; Figure 2B), as well as a time x sex interaction for both PO at GET (F_1,30_ = 6.93, *P =* 0.013, 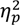 = 0.19) and PO at RCP (F _1,30_ = 4.19, *P =* 0.049, 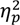 = 0.12). Indeed, PO at GET decreased only in males (-16 ± 15 %, *P =* 0.001) but not in females (-2 ± 13%, *P =* 0.460). Similarly, the decrease in PO at RCP was greater for males compared to females (-13 ± 10% vs -6 ± 9%, *P* = 0.049). A time effect was detected for V̇O_2_ at GET (F_1,59_ = 5.82, *P* = 0.019, 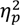 = 0.09; Figure 2C), with no sex effect or time x sex interaction (*P =* 0.096). No effects of time, sex, and time x sex interaction were found for V̇O_2_ at RCP (*P ≥* 0.169; Figure 2D).

**Figure 2.**
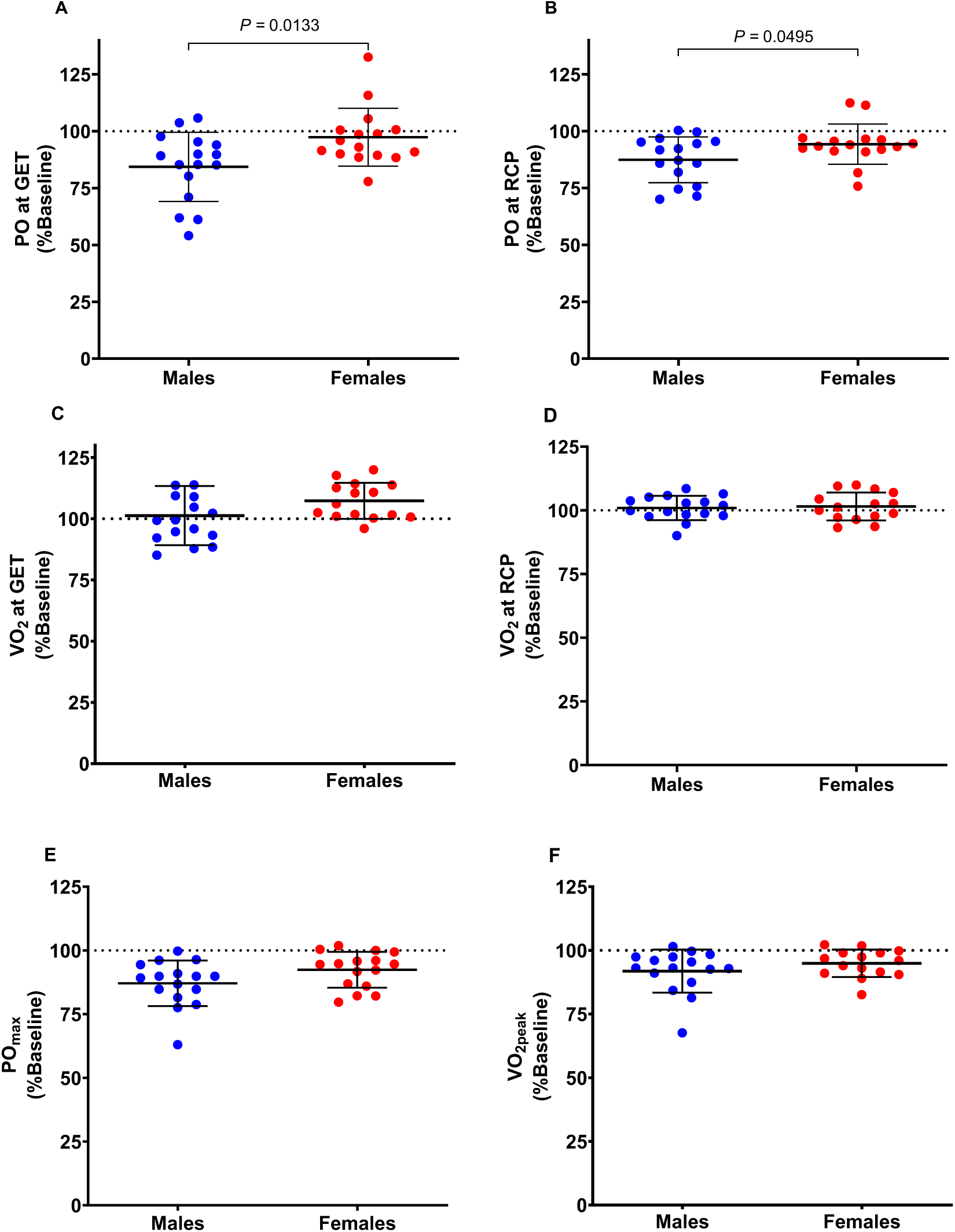
Incremental ramp test results after 90’ of heavy intensity cycling. A, power output at gas exchange threshold (GET); B, power output at respiratory compensation point (RCP); C, Oxygen uptake (V̇O2) at GET; D, Oxygen uptake (V̇O2) at RCP; E, maximal power output (POmax); F, peak oxygen uptake (V̇O2peak). Male data are presented in blue and female data in red. Dots indicate individual participants and lines indicate the group mean ± standard deviation.

### Responses during 90 mins

#### Pulmonary gas exchange

During HVY, a time effect was found for RPE (F _5,143_ = 51.06, *P* < 0.001, 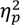 = 0.64), V̇E (%peak, F _6,176_ = 9.93, *P* < 0.001, 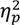 = 0.25), and fat oxidation (F _6,177_ = 11.95, *P* < 0.001, 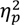 = 0.29) which all increased over time, as well as GE (F _6,175_ = 6.86, *P* < 0.001, 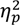 = 0.19), which decreased over time; all without main effects of sex or time x sex interactions (*P* ≥ 0.741). A time effect (F _6,168_ = 54.92, *P* < 0.001, 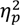 = 0.66) and time × sex interaction (F _6,168_ = 3.12, *P* < 0.006, 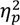 = 0.10) were detected for HR, demonstrating greater increases in males at 60 (*P =* 0.042) and 90 min (*P =* 0.032; Figure 3A). Similarly, RER demonstrated a time effect (F _6,175_ = 26.52, *P* < 0.001, 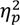 = 0.48;) and time x sex interaction (F _6,175_ = 3.84, *P =* 0.001, 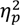 = 0.12), with greater decreases in females at 60 (*P =* 0.015) and 90 min (*P =* 0.016; Figure 3B). A sex effect was found for BLa (F _1,29_ = 5.24, *P* = 0.030, 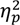 = 0.15) with greater blood lactate concentrations in males than females at 30 (*P =* 0.039), 75 (*P =* 0.022) and 90 min of exercise (*P =* 0.018; Figure 3C). A time effect (F _6,175_ = 17.96, *P* ≤ 0.001, 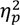 = 0.38), sex effect (F _1,30_ = 6.39, *P =* 0.017, 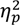 = 0.18) and time x sex interaction (F _6,175_ = 2.99, *P* = 0.008, 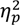 = 0.09) were found for V̇O_2_ (%peak), which increased more in males compared to females from 15min of exercise onwards (*P ≤* 0.021; Fig. 3D). Relative EE (time effect F _6,175_ = 16.70, *P* ≤ 0.001, 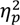 = 0.36; sex effect F _1,30_ = 21.01, *P* ≤ 0.001, 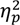 = 0.41; time x sex interactions F _6,175_ = 3.14, *P* = 0.006, 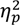 = 0.10) showed greater increases in males than females (*P ≤* 0.001; Fig. 4E). Whereas carbohydrate oxidation (time effect F _6,175_ = 4.02, *P =* 0.001, 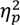 = 0.12; sex effect F _1,30_ = 41.62, *P* ≤ 0.001, 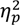 = 0.58; time x sex interactions F _6,175_ = 3.44, *P* = 0.003, 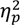 = 0.11) decreased in females from 45 min of exercise (*P ≤* 0.008) but not in males (*P ≥* 0.874).

**Figure 3.**
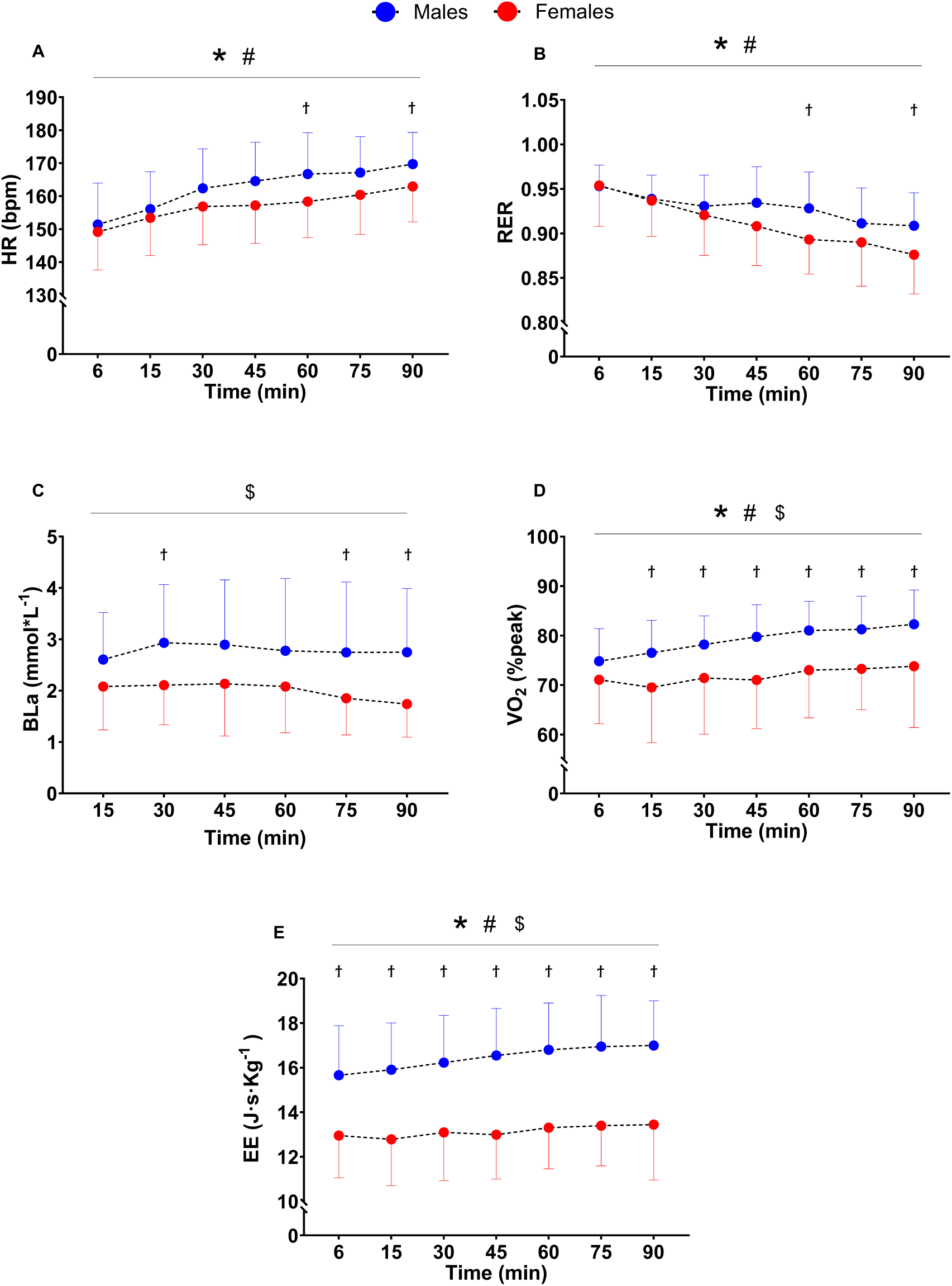
Physiological responses during 90’ minutes heavy intensity cycling. Changes in A, heart rate (HR); B, respiratory exchange ratio (RER); C, blood lactate (Bla); D, Oxygen uptake as %peak (V̇O2); E, energy expenditure (EE) during the 90’ constant heavy intensity cycling. Dots indicate group mean and lines indicate the standard deviation. *time effect; # time x sex interaction; $ sex effect; ϯ different between males and females.

#### Changes in neuromuscular function

The changes in neuromuscular function are displayed in Figure 4. A time effect was found for MVIC (-16 ± 12%; F_1,30_ = 62.57, *P* < 0.001, 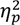 = 0.67; Figure 3A), Db_100_ (-15 ±13%; F_1,58_ = 40.51, *P* < 0.001, 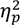 = 0.41), Db_10:100_ (-22 ± 11%; F_1,58_ = 135.9, *P* < 0.001, 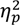 = 0.71), Q_tw·pot_ (-27 ± 13%; F _1,59_ = 132.80, *P* < 0.001, 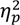 = 0.69), and VA (-6 ± 6%; F _1,59_ = 24.11, *P* < 0.001, 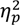 = 0.29; Figure 3B), without time x sex interaction effects (*P* ≥ 0.096). Although M_max_ did not show a main effect of time (*P* = 0.320), there was a sex effect (F _1,56_ = 11.99, *P =* 0.001, 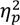 = 0.18) and a time x sex interaction (F _1,56_ = 11.99, *P =* 0.001, 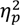 = 0.18), increasing in females (+19 ±35%, *P =* 0.003) and not in males (-12 ±21%, *P =* 0.082).

**Figure 4.**
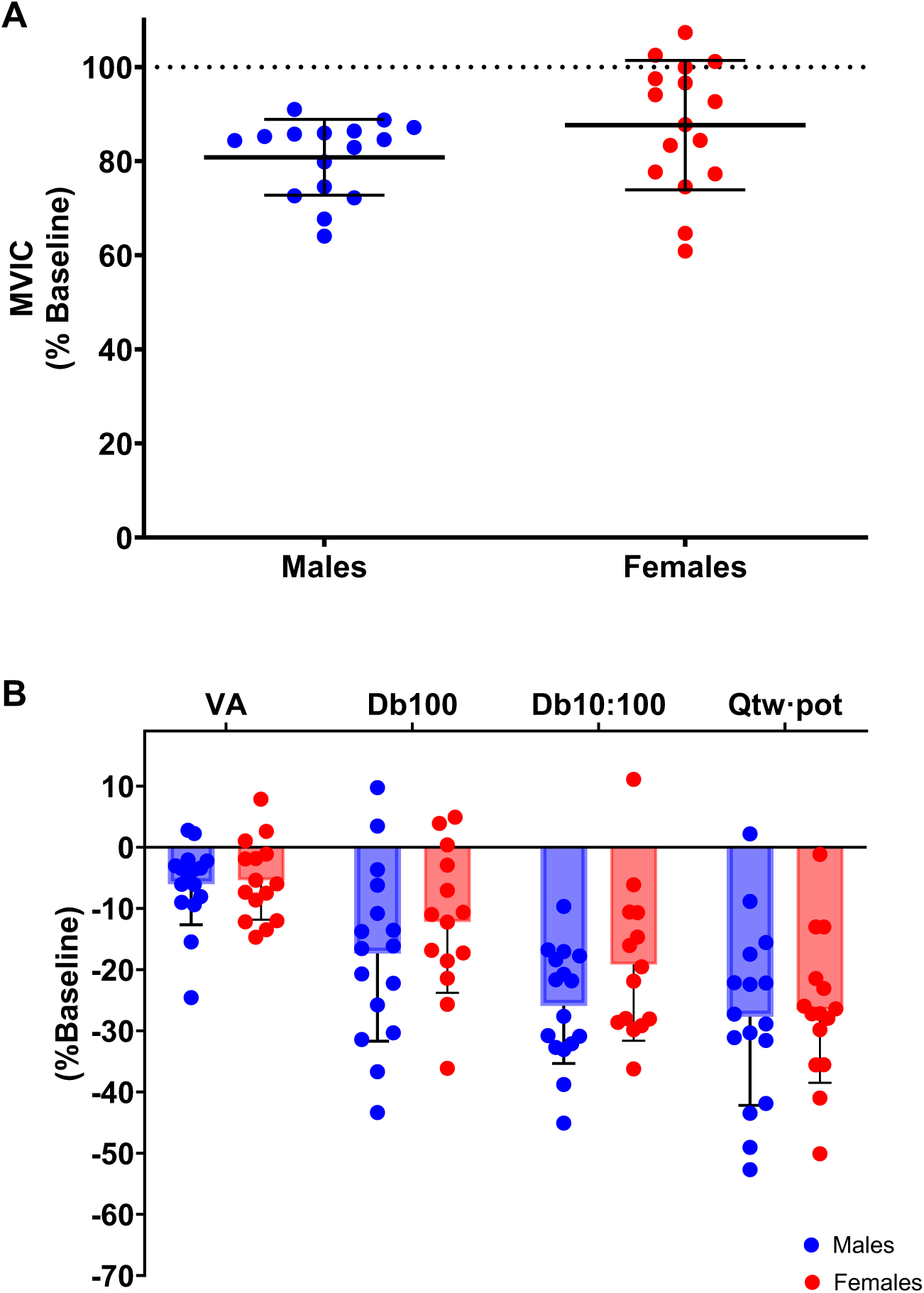
Changes in neuromuscular function. A, changes in maximal voluntary isometric contraction (MVIC); B, changes in potentiated twitches (Db100, Db10:100, Q_tw·pot_) and voluntary activation (VA) by sex from PRE to POST. Dots indicate individual participants and bars indicate the group mean ± SD. For all parameters, the PRE-POST decrease was statistically significant.

#### Electromyography

The EMG_RMS_/MVC during HVY did not change in the VL over time during the prolonged cycling task (time effects *P =* 0.560, initial values: 38.6 ± 18.6% MVC). Similar findings were reported for EMG_RMS_/M_max_ for VL (2.4 ± 1.4% M_max_, time effect *P ≥* 0.330).

#### Muscle oxygenation

A time effect (F _6,147_ = 22.99, *P <* 0.001, 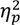 = 0.48) and time x sex interaction (F _6,147_ = 11.91, *P <* 0.001, 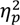 = 0.33) were found for O_2_HbMb (%ischemia) which increased in females but not in males from 45 min of exercise (Figure 5A). A time effect (F _6,147_ = 7.15, *P <* 0.001, 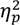 = 0.23) and time x sex interaction (F _6,147_ = 3.30, *P* = 0.005, 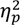 = 0.12) were found for HHbMb (%ischemia) which increased in males but not in females from 45 min of exercise (Figure 5B). A sex effect (F _1,24_ = 19.28, *P* = 0.001, 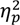 = 0.45) and time x sex interaction (F _6,139_ = 7.04, *P* < 0.001, 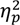 = 0.23) were found for TSI (%) which decreased more in males than females from 15 min of exercise (Figure 5C). Only a time effect (F _6,129_ = 17.62, *P* < 0.001, 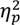 = 0.45) was found THb+Mb, which increased over time without sex or time x sex interaction effects (*P ≥* 0.079; Figure 5D).

**Figure 5:**
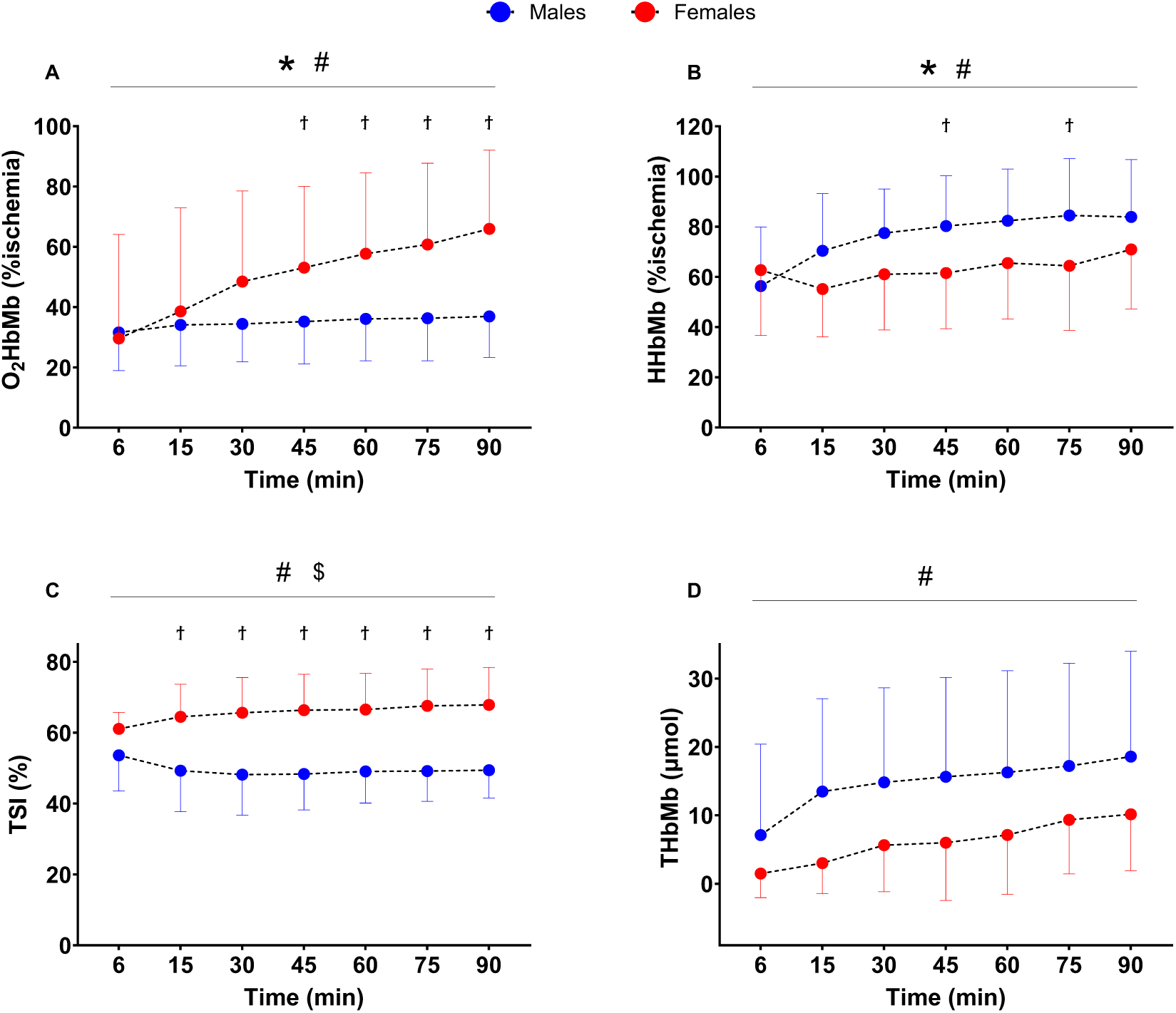
Indices of muscle oxygenation throughout the 90 minutes of heavy intensity cycling. A, oxyhaemoglobin and myoglobin (O_2_HbMb); B, deoxyhaemoglobin and myoglobin (HHbMb); C, tissue saturation index (TSI); D, total haemoglobin and myoglobin (THbMb). Dots indicate group mean and lines indicate the standard deviation. *time effect; # time x sex interaction; $ sex effect; ϯ different between males and females.

## DISCUSSION

The present study aimed to investigate durability and its physiological underpinnings in males and females following 90 minutes of constant-load cycling in the heavy intensity domain. Females exhibited lesser reductions in the power outputs associated with submaximal thresholds, whereas data recorded at peak exercise decreased similarly in both sexes. Surprisingly, no sex differences were observed in the changes in neuromuscular function, which was impaired similarly in males and females at the end of the fatiguing task. During the 90 minutes of heavy intensity cycling, heart rate, relative V̇O_2_ (%peak) and energy expenditure increased more in males than females, suggesting the development of a greater V̇O_2_ slow component and loss of efficiency as exercise proceeded. This was supported by the greater rise in HHbMb, suggesting elevated O_2_ extraction in males compared to females. Sex differences were also observed in substrate oxidation, with females demonstrating a lesser reliance on carbohydrates in the latter stages of exercise. These physiological sex differences observed during and following prolonged cycling exercise led to the conclusion that equivalently trained females have greater durability than their male counterparts.

### Incremental exercise test performance

After the 90 minutes of heavy intensity cycling, power output associated at GET decreased only in males by 16% whereas no significant decrease was observed in females. Similarly, the power output recorded at RCP decreased more in males (-13%) than females (-6%). The male decrement in the power output associated at RCP is in line with previous findings of Clark et al. (2018) and Clark at al. (2019a) who reported a reduction of the critical power by 8% and 11% respectively in males after 2 hours of cycling in the heavy intensity domain. However, the decrement of power output at GET was slightly greater than the 6-10% previously reported by Stevenson et al. (2022) and Hamilton et al. (2024) following moderate intensity cycling, which is likely explained by the difference in the intensity domain in which the participants performed the fatiguing cycling task, with the present study employing heavy intensity cycling. The higher intensity sustained by our participants could have led to a greater accumulation of fatigue throughout the cycling exercise (Brownstein et al. 2022), as suggested by the greater magnitude of change in in V̇O_2_, EE and RER, leading to a greater deterioration of performance during the ramp test performed in fatigued condition. This is not surprising, as recent scientific evidence (Leo et al. 2022; Spragg et al. 2024; Mateo March et al. 2024) reported that the intensity at which the fatiguing cycling task is performed is more crucial in determining the subsequent downward shift in the power duration relationship in professional cyclists rather than the total duration of the trial.

Although sex differences were observed at submaximal thresholds, values recorded at peak exercise, including P_max_ (-10 ± 8%), V̇O_2peak_ (-7 ± 7%), V̇CO_2peak_ (-15 ± 12%), and BLa_peak_ (-45 ± 20%) decreased similarly in males and females. These decreases in peak values are similar to those reported by Brownstein et al. (2022), who found a 6% reduction in V̇O_2peak_ and a 7% decrease in P_max_ after 90 minutes of cycling at 110% GET. Although the mechanism of P_max_ and V̇O_2peak_ decrease after a heavy intensity cycling exercise are not fully understood, several factors could have contributed. For example, participants experienced substantial impairments to excitation-contraction coupling (Ørtenblad et al. 2013), reducing their capability to express maximal power production (Clark et al. 2019a; Fuellerton et al. 2021). Indeed, the modest accumulation of metabolites that occurred during the fatiguing task could have contributed to a quicker attainment of critical levels of contractile dysfunction during the final stages of the incremental test (Black et al. 2017). Furthermore, the reduced V̇O_2peak_ could be attributed impairments in the O_2_ delivery to the working muscles associated with a decline in stroke volume (and consequently in cardiac output), as evidenced by the upward drift in HR during the 90-minute cycling, likely caused by an increase in core temperature, dehydration and reduced plasma volume (Nielsen et al. 1993). Alternatively, reductions in muscle activation at peak exercise could have contributed to reductions in O_2_ demands and V̇O_2peak_ given the reductions in VA found post-exercise in the present study. The greater reduction in submaximal markers of performance for males compared to females, without accompanying sex differences at peak exercise capacity, reveals insight into the potential mechanisms of the sex difference in durability. When these data are taken into consideration with those collected during the 90-minute fatiguing task (i.e. O_2_ uptake and extraction, substrate metabolism, and neuromuscular function) it is possible to gain insight into the etiology of the observed sex differences.

### Oxidative metabolism during and following heavy intensity exercise

In line with previous findings (Clark et al 2018, 2019a, 2019b; Stevenson et al. 2022; Hamilton et al. 2024), the 90 minutes of heavy intensity cycling trial resulted in a progressive deterioration of physiological parameters overtime. Indeed, HR, V̇O_2_, V̇E, EE increased, whereas GE and RER decreased as exercise proceeded, indicating that the accumulation of fatigue and physiological perturbation, due to both neuromuscular and metabolic changes, resulted in a loss of movement efficiency and impairment in performance. However, as reported above, females experienced a lower decline in submaximal thresholds, which is likely related to the lower deterioration of certain physiological parameters recorded during the 90-minute task.

At the beginning of the fatiguing exercise, the relative intensity of exercise (expressed as %HR_peak_ and %V̇O_2peak_) was similar between sexes (81% HR_peak_ and 73%V̇O_2peak_) indicating similar initial physiological demand. However, during the 90’ of fatiguing exercise, sex differences in the temporal change of both these two parameters appeared. Indeed, heart rate increased more in males than females from 60 minutes, as well as %V̇O_2peak,_ which increased more in males than females from 15 minutes onwards. The different trajectories of these parameters between sexes highlight the presence of a greater V̇O_2_ slow component in males (4.30 ml O_2_ · kg · min^-1^) compared to females (2.51 ml O_2_ · kg · min^-1^), which reached 82% V̇O_2peak_ in males and only 76% V̇O_2peak_ in females at the end of the trial (Figure 4D). This greater rise in V̇O_2_ was also accompanied by a larger increase in EE as well as in O_2_ extraction at the muscle level in males, as shown by their greater rise in HHbMb (+49%) than females (+13%) as exercise proceeded. There are multiple factors that could explain why this V̇O_2_ drift occurred, such as the V̇O_2_ slow component and/or a loss of mitochondrial efficiency. The mechanisms behind the V̇O_2_ slow component are not completely understood (Korzeniewski & Zoladz, 2015). However, it is reported to represent a progressive loss of skeletal muscle efficiency and homeostasis and therefore to be associated with the fatigue process (Jones et al. 2011). There is strong evidence that suggest that, during constant-work-rate exercise, the development of the V̇O_2_ slow component is associated with the fatigue-dependent increase ATP demand of the initially recruited motor units as well as the progressive recruitment of additional (type II) muscle fibres (VØllestad et al. 1984; Krustrup et al. 2008), which, in turn, determined a lower efficiency and a greater O_2_ cost (and thus V̇O_2_) of force production (Rossiter et al. 2002), potentially leading to a faster glycogen depletion, exacerbating the fatigue process. It is also well reported in literature that the V̇O_2_ slow component is greater in those individuals that possess a higher proportion of type II versus type I fibres in the vastus lateralis (Barstow et al. 1996; Pringle et al 2003; Jones et al, 2005). This evidence supports our findings, as females are well known to have a lower proportion and cross-sectional area of type II fibres compared to males (Nuzzo, 2023). Moreover, the greater contribution of glycolysis to ATP resynthesis and the increased metabolic instability withing the working muscles, as indicated by higher BLa values and increased HHbMb levels in males, without augmented muscle activation (i.e. EMG), could have contributed to a greater V̇O_2_ slow component in males (Colosio et al. 2020, 2021). It should be noted, however, that our data contradict previous findings whereby no sex difference in V̇O_2_ slow component or HHbMb increase were observed during heavy intensity cycling (Solleiro Pons et al. 2024), although the duration of exercise was far shorter (30 minutes) compared to the present study’s 90 minutes. Overall, our findings (i.e. the greater V̇O_2_ slow component, together with a larger increase in HHbMb in males), suggests that muscle contraction becomes less efficient in males, therefore requiring a higher energetic demand in order to maintain the same external power output.

The lesser decrease in efficiency demonstrated by females could also be related to sex differences in mitochondrial oxidative function (Cardinale et al. 2018). Indeed, a higher intrinsic mitochondrial respiration (i.e., the rate of mitochondrial respiration for a given amount of mitochondrial protein) as well as a lower mitochondrial O_2_ affinity were reported in females compared to males (Cardinale et al. 2018). This lower mitochondrial O_2_ affinity exhibited by females implicates a lower relative activation of mitochondria at similar relative intensity of exercise which is associated with higher aerobic mechanical efficiency during exercise (Larsen et al., 2011), lower ROS production, and better fat oxidation capacity (Montero et al. 2018). One further factor that could lead to greater energy expenditure and V̇O_2_ drift in males is the role of mitochondrial uncoupling, whereby a greater proportion of energy is dissipated as heat, rather than converted to ATP. Mitochondrial uncoupling occurs during prolonged periods of elevated oxygen consumption to protect against oxidative stress in response to increased ROS production (Brand & Esteves, 2005). As protons leak through uncoupling proteins, rather than generating ATP via ATP synthase, oxidative phosphorylation becomes less efficient, with more O_2_ consumption required to maintain a given rate of ATP production (Mogensen et al. 2006). Sex differences in ROS production during exercise have previously been observed, with oestrogen considered to limit this oxidative stress (for review see Enns & Tiidus, 2012), while evidence from rodents suggests that oestrogen supresses the expression of uncoupling protein 3 (Nagai et al. 2016). Therefore, although not directly studied in humans during exercise, there is the potential for female mitochondria to rely less on uncoupling to maintain homeostasis compared to males.

This greater loss of efficiency in males during the 90 minutes of heavy intensity cycling is a likely the primary contributing factor to the greater decreases in submaximal thresholds measured in the post-exercise incremental test. With no sex difference observed for the change in V̇O_2_ at GET or RCP, this implies that the change in metabolic rate at which the threshold occurred at was not different between sexes. However, the greater loss of efficiency in males meant that the metabolic rates at GET and RCP were attained at lower relative power outputs compared to females. These findings mirror those of Clark et al. (2019), who observed similar changes (i.e., a higher V̇O_2_ per watt of power output) with critical power following two hours of heavy intensity cycling; however, the present study extends this to females, and demonstrates that the loss of efficiency (and corresponding decrease in submaximal threshold power) is less severe compared to males.

### Substrate utilisation

Sex differences in the loss of efficiency could also be related to shifts in substrate metabolism during the 90-minute task. RER decreased more in females than males from 60 minutes of exercise, and at 45 min of exercise, carbohydrate oxidation dropped in females but not in males. This suggests that females relied more on fatty acid utilisation rather than glycogen to provide energy to the exercising muscles in the latter stages, which could elicit a glycogen-sparing effect. A lower reliance on carbohydrate oxidation in females is supported by evidence showing that males use 25% more muscle glycogen and have higher RER values than females at matched exercise intensities below CP (Tarnopolsky et al. 1990; Roepstorff et al. 2002). Indeed, as suggested by Roepstorff et al. (2006), the specific muscle morphology of females characterized by a higher proportion of type I muscle fibres and a greater capillarisation in the knee extensors could explain the higher fat oxidation and the improved muscle cellular energy balance (confirmed by the lack of AMPK activation in females during a fatiguing exercise at 60%V̇O_2peak_) exhibited by females during prolonged endurance exercise. This could potentially lead to a better preservation of glycogen stores overtime and preserve performance, as glycogen depletion is an important limiting factor of prolonged exercise performance (Bergstrom et al. 1967; Hermansen et al. 1967; Ørtenblad et al. 2013). It should be noted that in our study, the lesser decrease in RER, together with a maintained carbohydrate oxidation in males compared to females, was likely also a result of the higher relative energy demand (i.e., % V̇O_2peak_) as exercise proceeded, in order to maintain the target power output.

Moreover, to further support this sex differences in metabolism, in our study males showed greater blood lactate concentrations at each time point of the 90’ of heavy intensity cycling. However, blood lactate did not increase progressively through the task in either group, indicating that exercise was performed in a steady state in the heavy intensity domain. The higher blood lactate concentration that was found in males than females could be explained by sex differences in muscle metabolism (Devries, 2015). Similarly to our findings, it is well reported in literature that during endurance exercise males rely more on carbohydrate oxidation to support fuel requirements and they display a greater activation of the glycolytic metabolic pathway compared to females, which is probably related to sex differences in fibre type and anaerobic metabolic properties of skeletal muscle (Esbjornsson et al. 1993). This greater glycogen preservation in females could conceivably contribute to the lesser reductions in the power outputs at metabolic thresholds that were observed after the 90-minute task. Indeed, in states of glycogen depletion, V̇O_2_ per unit of power output is increased due to the lesser ATP yield per oxygen consumed during fat metabolism (Krustrup et al. 2004). Therefore, in addition to oxidative differences between sexes, differences in substrate metabolism likely contributed to the observed sex difference in durability.

### Neuromuscular function

The decrement in voluntary force (-16% in MVIC) after the 90 minutes of heavy intensity cycling was a result of both central and peripheral adjustments. However, both decrements in voluntary activation as well as in resting potentiated twitches were reported without sex differences. Similarly, RPE increased in both sexes without differences. The lack of sex differences in neuromuscular fatigue contradicts previous literature showing lower levels of fatigability in females compared to males during various cycling tasks (Glace et al. 2013; Ansdell et al. 2020; Azevedo et al. 2021). The reasons behind this discrepancy are unclear. However, as reported by Hunter (2016), while there is a general consensus that females are less fatigable than males during and after intermittent and sustained submaximal isometric contractions, data concerning whole-body dynamic contractions are more equivocal, and dependent on the task performed. Previously, it has been suggested that neuromuscular fatigue could contribute to the reduction in submaximal thresholds (Clark et al. 2019). Our data does not necessarily refute this, but rather it suggests that sex differences in durability are likely not due to sex differences in neuromuscular fatigue resistance. Instead, as mentioned above, sex differences in oxidative function and substrate metabolism likely play a larger role.

### Experimental Considerations

As expected, males showed higher absolute and relative V̇O_2peak_, with the latter being 12% greater in males than females. These results are not surprising, as the sex difference in relative V̇O_2peak_ is reported to be approximately 10-14% greater in males than females among similarly trained cohorts (Pate & O’Neill, 2007). As reported by Joyner (2017) the higher V̇O_2peak_ values in males compared to females may be primarily due to a higher cardiac output, haemoglobin concentrations, and lesser fat mass exhibited by males. Therefore, the sex differences in durability and physiological derangements during and following heavy intensity cycling are likely not due to sex differences in training status.

Finally, the females in the present study were tested in a low endogenous hormonal state. Although the role of sex hormones on factors such as metabolic thresholds (James et al. 2023), substrate metabolism (D’Souza et al. 2023) and oxidative function (Mattu et al. 2020) during exercise are thought to be minimal, it remains to be determined whether durability is altered by factors such as the menstrual cycle and contraceptive usage. This knowledge is crucial for female athletes and those working with them, allowing practitioners to confidently base their prescription on data generated in females.

## CONCLUSION

This study demonstrated that after 90 minutes of heavy intensity cycling, females experienced lower reductions in the power outputs at submaximal thresholds than males, confirming our hypothesis of superior durability in females. In contrast, maximal exercise capacity was impaired to a similar degree in both sexes. No sex differences were observed in the changes in neuromuscular function, which was impaired similarly at the end of the fatiguing task. However, during the 90-minute task, females demonstrated lesser V̇O_2_ drift and rise in muscle oxygen extraction, a progressively greater reliance on fat oxidation, lower blood lactate values, as well as a greater muscle oxygenation. These results suggest that the better preservation of performance exhibited by females in our study is not directly linked to sex differences in neuromuscular function. However, sex differences in substrate utilisation and oxidative metabolism seem to play a more important role in delaying the loss of efficiency. The better preservation of submaximal thresholds in females after the prolonged fatiguing task highlight the need to consider sex-specific training prescription and pacing strategies for long duration events.

## Additional information

### Data availability statement

The data that support the findings of this study are available from the corresponding author upon reasonable request.

### Autor contributions

E.P., M.C., S.P., C.B and P.A. contributed to the conception and design of the work; E.P., P.S., E.S., L.B., H.K.W., M.C., C.F., A.M., S.C., B.M., W.P. and C.B. collected the data; E.P., P.S., E.S., P.S., M.C., C.F., A.M., S.C., B.M., W.P. and C.B analysed the data; E.P., M.C., S.P., C.B. and P.A. interpreted the data; E.P. drafted the article; E.P., E.S, P.S., M.C., S.P., C.B and P.A revised the article. All authors approved the final version of the article, agree to be accountable for all aspects of the work in ensuring that questions related to the accuracy or integrity of any part of the work are appropriately investigated and resolved. All persons designated as authors qualify for authorship, and all those who qualify for authorship are listed.

## Acknowledgements

The researchers would like to thank the participants of the present study for their time and efforts and Stefano Chiaverini, a master student from the University of Pavia who helped with data collection.

